# Aging-related olfactory loss is associated with olfactory stem cell transcriptional alterations in humans

**DOI:** 10.1101/2021.08.09.455538

**Authors:** Allison D. Oliva, Khalil Issa, Ralph Abi Hachem, David Jang, Rupali Gupta, E. Ashley Moseman, Hiroaki Matsunami, Bradley J. Goldstein

## Abstract

Presbyosmia, or aging related olfactory loss, occurs in a majority of humans over age 65 years, yet remains poorly understood, with no specific treatment options. The olfactory epithelium (OE) in the nasal fossa is the peripheral organ for olfaction, and is subject to acquired damage, suggesting a likely site of pathology in aging. OE basal stem cells reconstitute the neuroepithelium in response to cell loss under normal conditions. In aged OE, patches of respiratory-like metaplasia have been observed histologically, consistent with a failure in normal neuroepithelial homeostasis or repair. Accordingly, we have focused on identifying cellular and molecular changes in presbyosmic OE. Combining psychophysical testing with olfactory mucosa biopsy analysis, single cell RNA-sequencing (scRNA-seq), and human olfactory culture studies, we identified evidence for inflammation-associated changes in the OE stem cells of presbyosmic patients. The presbyosmic basal stem cells exhibited increased expression of genes involved in response to cytokines or stress, or the regulation of proliferation and differentiation. To facilitate further study of human OE stem cells, we developed an adult human basal cell culture model. Characterization of cultures using scRNA-seq confirmed maintenance of a reserve stem cell-like phenotype, and brief cytokine exposure in basal cell cultures resulted in increased expression of TP63, a transcription factor acting to prevent OE stem cell differentiation. Our data are consistent with a process by which aging-related inflammatory changes in OE stem cells may contribute to presbyosmia, via the disruption of normal epithelial homeostasis, suggesting that OE stem cells may represent a rational therapeutic target for restoration of olfaction.

**One Sentence Summary:** Single cell profiling suggests that inflammatory-associated olfactory epithelial stem cell dysfunction is associated with presbyosmia in humans.

## INTRODUCTION

The olfactory system detects volatile odors, providing chemosensory input via the first Cranial Nerve. Olfaction contributes to daily social interactions and quality of life, is important in the detection of smoke or dangerous chemicals, and underlies flavor perception in combination with gustatory input from taste buds. Loss of olfaction can result in depression, nutritional disorders, and increased mortality (*1-3*). Acquired olfactory loss, termed anosmia, can occur due to trauma, viral infections including COVID19, or sinonasal disease such as sinusitis (*4-6*). In addition, age-related loss of smell (presbyosmia) has been well described but remains poorly understood. Population studies using psychophysical testing have identified that half of subjects over age 65 years and greater than two-thirds of those over the age of 80 exhibit impaired olfaction (*7, 8*). However, there are no effective therapies for presbyosmia. To facilitate development of specific treatments, an understanding of the mechanisms underlying this condition is required.

Limited to the olfactory cleft, the OE is a true neuroepithelium containing bipolar chemosensory neurons, while the remainder of the nasal cavity is lined by respiratory epithelium consisting of ciliated and secretory cells. The OE contains neurogenic populations known as globose basal cells (GBCs) and horizontal basal cells (HBCs) (*9*). In rodent models, during normal olfactory sensory neuron turnover the GBCs divide and differentiate to produce new neurons while the HBCs are relatively quiescent (*10-12*). However, when faced with injuries that disrupt the sustentacular cell layer of the OE, the reserve HBC stem cells are activated and serve as the main source of olfactory sensory neurogenesis (*13-16*). Human OE is exposed continuously to potential damage in the form of viral or other infections, inflammation driven by allergens and other inspired irritants, or trauma. Thus, the proper function of the neurogenic cycle is critical for homeostasis within the OE and, in turn, a functioning sense of smell.

Consistent with rodent models, neurogenic basal cells are active in the OE of adult humans (*17, 18*). Thus, one potential explanation for presbyosmia is an aging-related process of neurogenic exhaustion. In this model, aged adult neurogenic cell populations are believed to be exhausted and either no longer present or, if present, incapable of reconstituting the neuronal populations (*19*). As a result, as neurons are depleted with normal ongoing damage or “wear-and-tear”, the ability to maintain an intact OE is lost. Histologic study of olfactory biopsies or autopsy samples demonstrating areas of respiratory-like metaplasia in aged adults (*18-20*) supports the idea that the neuronal layer can be depleted, leaving an aneuronal, respiratory-like barrier epithelium in its place (Fig. 1A, 1B). However, adult olfactory neurogenesis is a highly regulated process involving a complex cellular milieu including immune cells, supporting cell populations, ensheathing glia, stromal and vascular cells, and multiple populations of basal stem and progenitor pools. While histologic findings have been described, direct evidence for molecular alterations in neurogenic OE populations in presbyosmic humans is lacking, limiting the ability to identify new treatment strategies for aging-related olfactory loss.

**Fig. 1.**
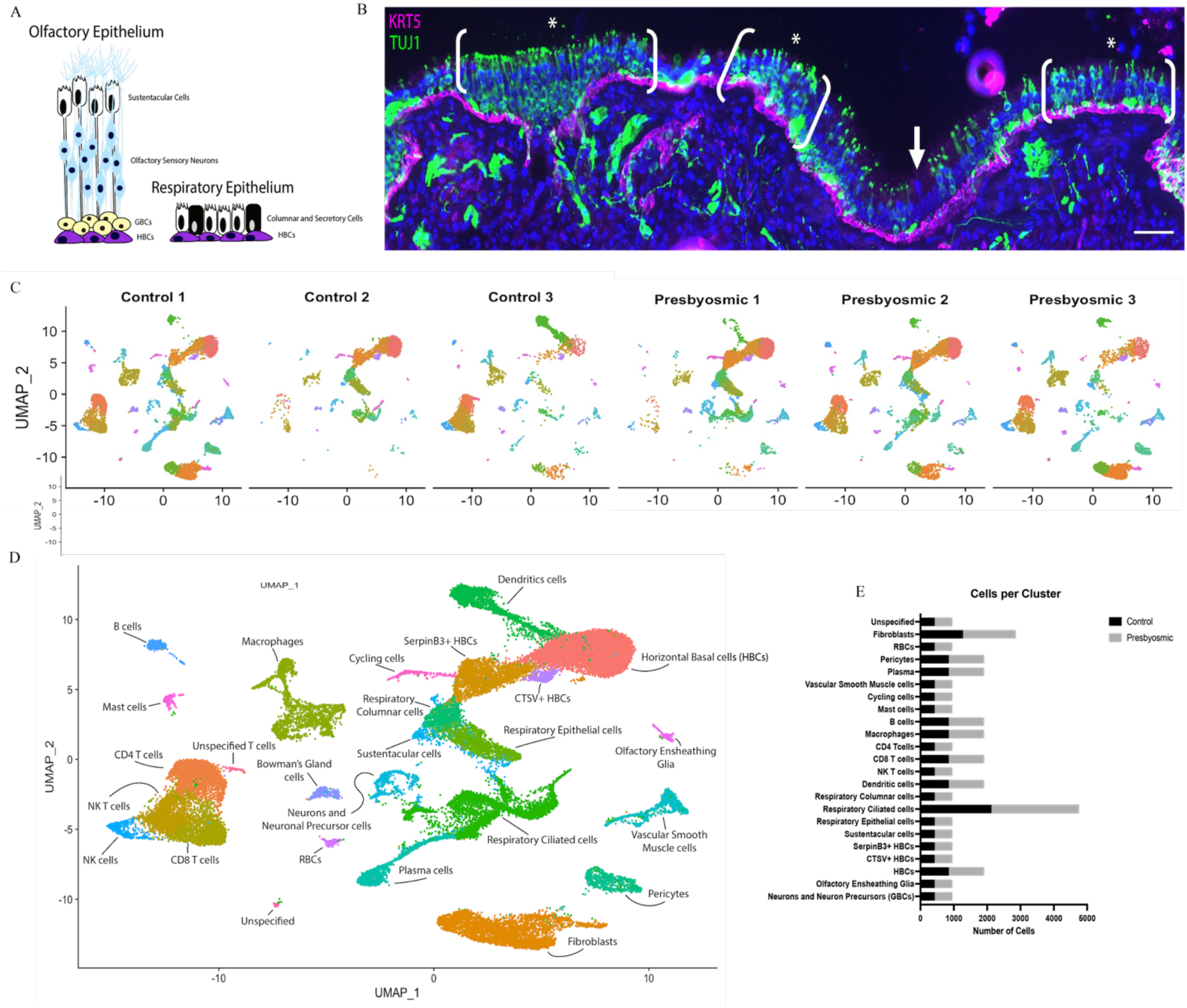
Integrated analysis of 36,091 cells from human olfactory epithelial biopsies of normosmic and presbyosmic adults. **(A)** Schematic illustration of cell types in human olfactory and respiratory nasal epithelium. HBCs = Horizontal basal stem cells; GBCs = Globose basal stem cells. **(B)** Human OE from an 80yo presbyosmic subject; brackets and asterisks indicate areas containing TUJ1+ OE neurons; a patch of aneuronal respiratory-like metaplasia is marked by the white arrow; HBCs are labeled with anti-KRT5. **(C)** UMAP projection showing integrated dataset split into separate projections by sample. **(D)** UMAP projection of combined dataset; clusters are annotated according to canonical markers. **(E)** Cell contributions from normosmic or presbyosmic cohorts, per cluster. Scale bar = 50 μM in B; nuclei are stained with DAPI (blue) in B.

Classic fate mapping studies following rodent OE damage defined HBCs as the multipotent OE reserve stem cell, activated by severe injury (*13*). Of interest, an experimental model of severe chronic olfactory inflammation in mice, involving OE exposure to TNFα, identified that HBCs may adopt a protective “barrier defense” transcriptional program, driven by NFKB or JNK signaling (*21*). In this model, prolonged inflammation prevents stem cell neurodifferentiation, despite neuron depletion and increased signaling between HBCs and immune cells (*22*). Together, the data from rodent olfactory injury-regeneration studies, and the potential influence of inflammatory signaling, suggest a model in which some human olfactory disorders, such as presbyosmia, may involve OE stem cell derangements linked to local microenvironmental cues.

Accordingly, the aim of this study is to investigate transcriptional changes in presbyosmic adult human OE samples at single cell resolution. The single cell approach is useful for identifying unbiased gene expression patterns in highly heterogenous tissues and enabling differential gene expression analysis between subpopulations of interest. We hypothesized that olfactory basal stem cells in older adults with hyposmia exhibit gene expression changes in pathways involved with neurogenic activity, compared to normosmic controls. To test this hypothesis, we performed scRNA-seq analysis of human OE biopsies from normosmic control subjects and hyposmic subjects aged 65 years or older, as well as culture and immunohistochemical assays. All subjects included in our study underwent psychophysical olfactory function testing using the Smell Identification Test (SIT) to measure their olfactory ability.

## RESULTS

### Study cohort

To investigate mechanisms involved with presbyosmia, we have obtained olfactory mucosa biopsies from 8 adult human subjects, ages 51 - 80 years, 4 male and 4 female (demographic details provided in **Table 1**). We sought controls with normosmic function, and therefore included one subject slightly below age 65 to meet these criteria, but otherwise avoided inclusion of young samples. Smell Identification Test (SIT) scores >33 are considered normosmic, 30-33 mild hyposmic, 26-29 moderate hyposmic, 19-25 severe hyposmia, 6-18 anosmic (with slight adjustments for age/gender). We processed samples for scRNA-seq from 6 subjects. One cohort consisted of 3 normosmic subjects, based on psychophysical olfactory assessment, with SIT scores 34 – 37; the other cohort consisted of presbyosmic subjects, SIT scores 11 - 29 and age >65 years. Active rhinitis or sinusitis was excluded by nasal endoscopy and/or non-contrast sinus CT scan showing absence of nasal polyps, severe edema or purulence. The diagnosis of presbyosmia was established based on SIT score and absence of other obvious cause for sensorineural hyposmia, such as prior head trauma, post-viral olfactory disorder, or neurodegenerative disease. Biopsies were obtained from the olfactory cleft at the time of transnasal endoscopic surgery for access to the sphenoid or skull base.

**Table 1.** Human subject demographics.

| Age | SIT score | Cohort | Use | Gender | Race |
| --- | --- | --- | --- | --- | --- |
| 71 | 37 | Control1 | scRNA-seq | Female | White |
| 66 | 34 | Control2 | scRNA-seq | Female | White |
| 51 | 37 | Control3 | scRNA-seq | Male | White |
| 78 | 29 | Presbyosmic1 | scRNA-seq | Male | Black |
| 74 | 11 | Presbyosmic2 | scRNA-seq | Female | White |
| 73 | 27 | Presbyosmic3 | scRNA-seq | Female | White |
| 69 | 24 | Presbyosmic | Histology | Male | Black |
| 80 | 28 | Presbyosmic | Histology | Male | White |

### Epithelial Analysis

For olfactory mucosa biopsies, we generated high quality single cell transcriptional profiles for 36,091 cells. Despite being obtained from the superior posterior septum, typically lined by olfactory surface epithelium (OE) rather than sinonasal respiratory surface epithelium (RE), biopsies were comprised by mixtures of both OE and RE cells (**Fig. 1 A-C**). Mixed epithelial populations are not unexpected, since there is a well-documented patchy replacement of OE by respiratory-like surface epithelium in adult humans (*18, 20*). An example of discontinuous intact neuroepithelium with aneuronal patches is shown in a presbyosmic biopsy stained to visualize olfactory neurons with antibody Tuj1, recognizing neuron-specific beta-tubulin (**Fig. 1B**). Using unsupervised clustering of scRNA-seq data, we visualized cellular composition in uniform manifold approximation projection (UMAP) plots; distinct populations were annotated as we and others have described, based on known marker gene expression (*17, 23, 24*). Clustering was visualized for individual biopsies (**Fig. 1C**) and in an integrated and batch-corrected combined plot (**Fig. 1D**). Because we used full-thickness surgical punch-style biopsies rather than a brush biopsy technique, our samples captured surface epithelial cells as well as large numbers of stromal cells, vascular cells and immune populations (**Fig. 1D**). While one presbyosmic subject with total anosmia had nearly no olfactory neurons present, all other samples contained sensory cell populations, ranging from 63 to 186 cells per sample (**Fig. 1C, 1D and Supplemental Fig.1**). Neuron lineage cluster analysis identified GBCs, immature and mature olfactory neurons, and identified expression of olfactory receptor transcripts, 56 of which were not found in our previous scRNA-seq data set (**Supplemental Fig.1)**. The relative cellular composition is indicated for presbyosmic and normosmic cohorts (**Fig. 1D**). DotPlot visualization of transcripts highly enriched in specific cell clusters provides an overview of annotated cell phenotypes (**Supplemental Fig.2**). Initial analysis demonstrates that our samples capture the range of cell populations present in olfactory mucosa. While this approach provides limited ability to draw definitive conclusions about relative numbers of specific cell populations, single cell profiling is particularly well suited to explore transcriptional changes focused on identical cell populations from diseased versus control cohorts.

### Focused cell cluster analysis: olfactory stem cells

Because a patchy replacement of neuroepithelium with respiratory-like metaplastic surface epithelium, lacking neurons, is a prominent feature of presbyosmic biopsies (**Fig. 1B**), a process of “neurogenic exhaustion” has been proposed as an underlying mechanism (*25*). In this model, olfactory basal stem or progenitor cell populations would exhaust or change over time, resulting in an inability to support ongoing epithelial renewal. Accordingly, we focused attention to the stem cell clusters in presbyosmic versus control normosmic samples (**Fig. 2**). The HBC, defined by KRT5 and TP63 expression, functions as the reserve stem cell population in the OE (*13*), and a biochemically similar basal cell also supports self-renewal of the respiratory epithelium (*24*). Re-plotting the KRT5^+^ basal cell cluster subset and visualizing sample contributions, we found that olfactory stem cells (HBCs) are abundant in presbyosmic biopsies (**Fig. 2A, B**). Nine sub-clusters were generated from >4000 cells, with contributions from all biopsy samples (**Fig. 2A**).

**Fig. 2.**
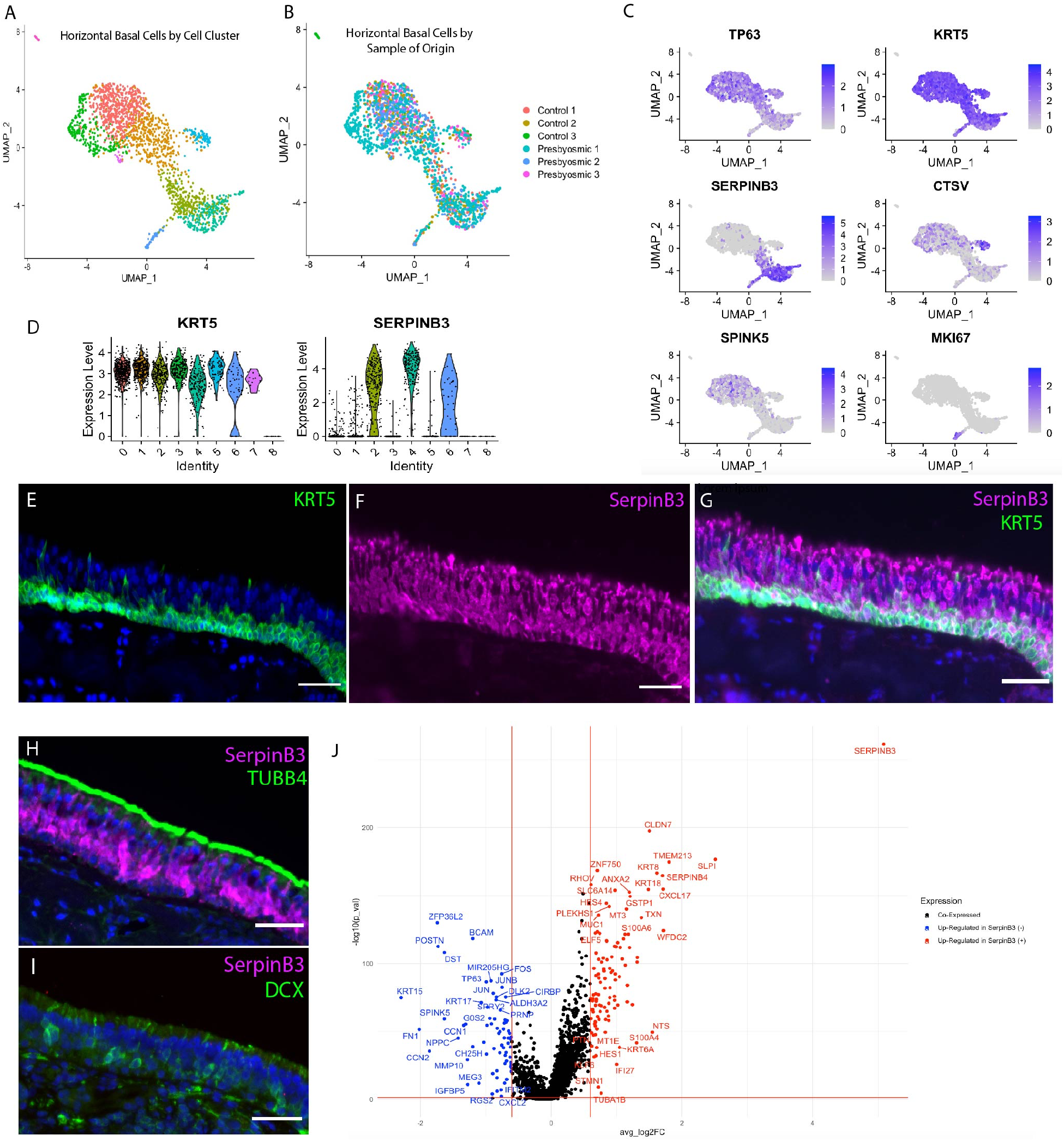
Horizontal basal stem cell transcriptional heterogeneity. **(A)** UMAP plot visualizing the HBC subset from integrated dataset after unbiased re-clustering. **(B)** HBC subset identifying cells by sample of origin. **(C)** Selected gene expression plots demonstrating heterogeneity among HBCs. **(D)** Comparison of KRT5 and SERPINB3 expression across HBC sub-clusters. **(E-G)** Human respiratory epithelium immunohistochemistry with SERPINB3 (magenta) expression in KRT5 (+) basal cells and the columnar or secretory cells. **(H)** Respiratory epithelium with co-localization of TUBB4 protein (green) apically and SERPINB3 (magenta). **(I)** OE from a normosmic subject co-stained for immature neuronal marker DCX (green) and SERPINB3 (magenta), absent. **(J)** Differential expression between the SERPINB3 (+) and SERPINB3 (-) HBCs. Genes significantly upregulated (p < 0.05 and log fold change >0.60) in SERPINB3 (+) cells are colored in red; genes significantly upregulated in SERPINB3 (-) cells are colored in blue. Nuclei in (E-I) are stained with DAPI (blue); bar = 50 μM.

### HBC heterogeneity

We identified here heterogeneity among the TP63^+^/KRT5^+^ basal cells (**Fig. 2C**). For instance, SERPINB3 strongly segregates to a subset of the basal cells present in our samples, as visualized in FeaturePlots or violin plots (**Fig. 2C, D**). In our prior scRNA-seq data set, SERPINB3 appeared to be enriched in respiratory rather than olfactory basal cells (*17*); here, immunohistochemical staining confirms at the protein level that SERPINB3 co-localizes in many KRT5+ basal cells in portions of surface epithelium *lacking* olfactory neurons but co-expressing known respiratory epithelial markers such as TUBB4 (also referred to as β4-tubulin (*25*)) (**Fig. 2E-I**). SERPINB3 is also present in more mature respiratory secretory cells (**Fig. 2F,G)**. However, observed areas harboring olfactory neurons, labeled with neuron-specific markers such as DCX, lack SERPINB3 (**Fig. 2I**). Thus, SERPINB3 may be useful in distinguishing olfactory from respiratory cells in heterogenous samples. Differential gene expression (DE) analysis comparing SERPINB3(+) versus SERPINB3 (-) basal cells identified several genes that are significantly (p < 0.05, log2FC > 0.60) upregulated in SERPINB3 (+) basal cells (**Fig. 2J**). Some of these enriched genes, such as HES4 and MUC1, have been demonstrated in respiratory cell lineages (*24*), and genes such as KRT8 and KRT18 are established non-neuronal markers in murine models (*26*). In contrast, transcripts that were enriched in SERPINB3 (-) HBCs including KRT15, MEG3, CTSV and SPINK5 are less well-studied. Visualizing SPINK5 and CTSV in UMAP plots, both transcripts localized broadly among the putative olfactory HBC clusters, although a small highly CTSV-enriched subset is identifiable (**Fig. 2C**). CTSV is of potential interest, as it is a druggable cysteine protease with restricted CNS expression, thought to have roles in regulation of cell proliferation, adhesion and, importantly, immune response (*27, 28*). SPINK5 encodes a proteinase inhibitor, also present in epidermal basal layers, with activity regulating proliferation via the Wnt pathway (*29, 30*); a role in HBCs has not been described.

### Presbyosmic versus control stem cells

We also performed DE analysis comparing HBCs from presbyosmic versus control groups (**Fig. 3**). Our analysis identified 31 transcripts significantly upregulated in presbyosmic HBCs compared to normosmic control HBCs, including cytokine-response genes IFI6, IFI27, IRF1, KLF4, KLF6, and anti-proliferative gene RHOB (**Fig. 3A**) (*31-33*). Gene set enrichment analysis (GSEA) of the upregulated set suggested strongly that the presbyosmic HBCs exhibit changes consistent with chronic inflammation. GSEA biological process categories included processes “cellular response to cytokine stimulus”, “regulation of cell proliferation”, “regulation of cell differentiation”, and numerous terms relating to regulation of apoptosis or cell death (**Fig. 3B**). Of interest, our findings from presbyosmic human HBCs are consistent with basal cell alterations identified in a mouse model of chronic local cytokine overexpression (*22*), providing support for a model of stem cell dysfunction in aging-related human olfactory loss.

**Fig. 3.**
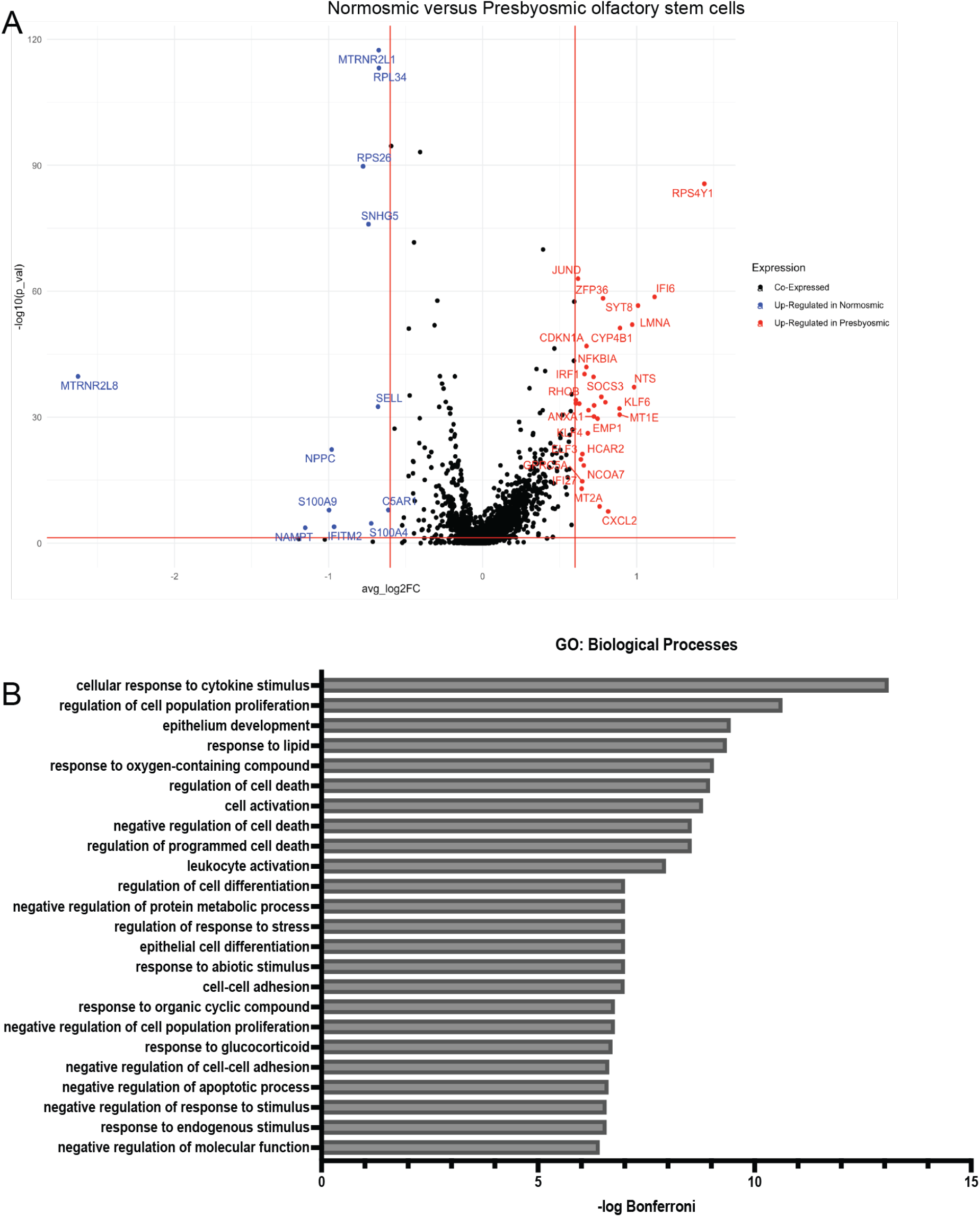
Differential gene expression identifies presbyosmic stem cell dysfunction. **(A)** DE analysis between presbyosmic and normosmic HBCs; transcripts significantly upregulated (p < 0.05 and log fold change >0.60) in presbyosmic HBCs (red), and transcripts upregulated in normosmic HBCs (blue) are labeled. **(B)** Gene set enrichment analysis of presbyosmic HBC-enriched transcripts, plotting the top significant biological process terms; inflammatory or injury response terms suggest HBC functional alterations. GO=gene ontology.

### Immune landscape in presbyosmic olfactory mucosa

Because we identified changes suggesting inflammatory responses in presbyosmic HBCs, we were interested in the immune cells captured in olfactory biopsies. Focusing attention to immune cell populations within our integrated scRNA-seq dataset, we investigated differential gene expression between the presbyosmic and normosmic samples. CD8^+^ T cells, CD4^+^ T cells, innate lymphocytes (NK, NKT and innate lymphoid cells (ILCs), CD68^+^ macrophages, and mast cells were identified (**Supplemental Fig. 3**). Initial analysis suggested possible differences among ILCs, so this population was further analyzed. The innate lymphocyte compartment (NK/ILC) cells from presbyosmic and normosmic control biopsies were re-clustered and visualized (**Fig. 4A, 4B**). While canonical “NK” markers, NKG7 and GNLY, were expressed uniformly (**Fig. 4C**), DE analysis identified several upregulated genes in cells from presbyosmic subjects (**Fig. 4D**). The most significantly upregulated gene was amphiregulin (AREG). AREG has been found to increase regulatory T cell proliferation, and to protect and maintain epithelial stem cells in corneal epithelial injury models (*34, 35*). But perhaps most intriguingly, a subset of innate lymphoid cells, ILC2s, are known to secrete AREG to bolster mucosal epithelial repair. AREG is a ligand for the EGF receptor, which is strongly expressed by olfactory HBCs, providing a mechanism of action to influence presbyosmic HBC activity (*34*). Indeed, ILC2 derived amphiregulin has been shown to play a prominent role in lung epithelial regeneration (*36*). Other genes upregulated in presbyosmic patients (**Fig. 4D**), including H3F3B, SYTL3 and HOPX are also associated with ILC2s (*37, 38*). Likewise, we identified the C-C motif chemokine ligand CCL3 and CCL4 as enriched in presbyosmic cells, and chemokine signaling between immune cells and HBCs is described in rodent models (*22*). Both CCL3 and CCL4 are reportedly expressed by sub groups of ILCs suggesting a potential influx or differentiation of multiple ILC subsets, or one plastic subset (*37*). These data point to potential mechanisms for ILC derived pro-epithelial signals to influence olfactory versus respiratory replacement within olfactory cleft.

**Fig. 4.**
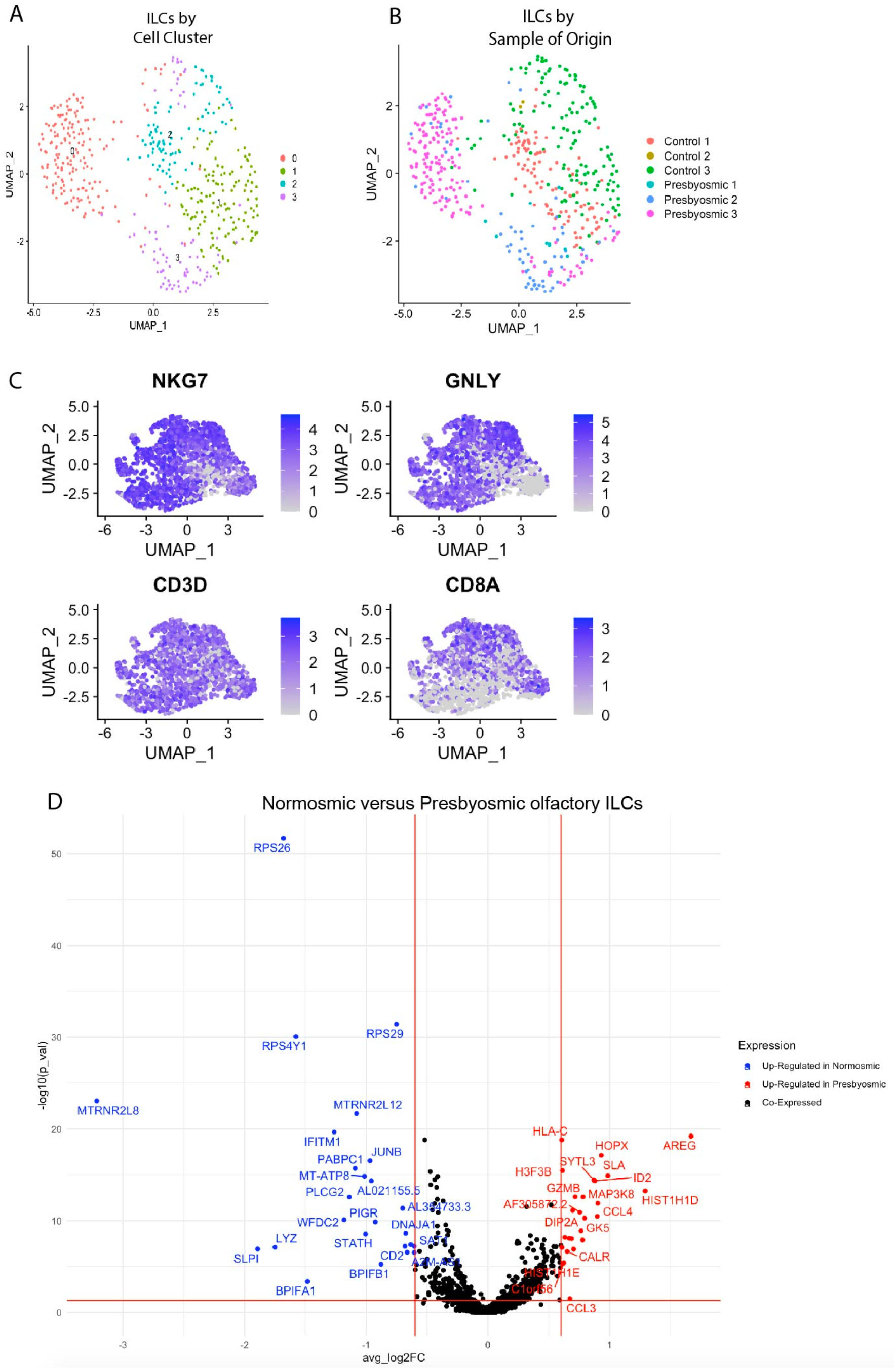
Gene expression changes in presbyosmic immune cells. **(A)** A UMAP plot of the subset of innate lymphocyte compartment cells after re-clustering. **(B)** sample origins are depicted. **(C)** Gene expression plots for selected innate lymphocyte compartment (NK/ILC) transcripts, (NKG7, GNLY, CD3D and CD8A); additional markers are depicted in Supplemental Fig. 3. **(D)** Differential gene expression between presbyosmic and normosmic ILCs; transcripts significantly upregulated (p < 0.05 and log fold change >0.60) in presbyosmic cells are colored in red, while transcripts significantly upregulated in normosmic are colored in blue.

### HBC culture model

To facilitate further mechanistic study of human olfactory stem cells, we sought to develop a human olfactory HBC culture model. We reasoned that immunoselection for cells expressing NOTCH1 from cell suspensions of human olfactory mucosa biopsies would enrich samples highly for HBCs, as NOTCH1 surface receptor has been demonstrated on HBCs (Herrick et. al., 2017). Thus, we prepared a surgical olfactory mucosal biopsy sample from a normosmic subject for scRNA-seq immediately following tissue harvest, and the remaining cell suspension was then NOTCH1 immunoselected for culture (**Fig. 5A**). The NOTCH1 (+) fraction was maintained on 5% laminin in a well-established respiratory basal cell growth medium for several passages. Importantly, the scRNA-seq analysis of the portion of the biopsy used to establish this culture revealed almost exclusively olfactory rather than respiratory populations (see **Fig. 1C, 1D**). Thus, we conclude that the NOTCH1+ cells seeded to establish our cultures were likely to be olfactory HBCs, rather than possible respiratory epithelial contaminants. The cultured cells consistently formed adherent islands with occasional morphological changes consistent with differentiation (**Fig. 5 B**). Representative immunocytochemical staining (**Fig. 5 C**) showed nearly homogenous expression of HBC markers KRT5 and TP63. Passaged cultures were further characterized using scRNA-seq (**Fig. 5 D, E**). Integrating and visualizing the scRNA-seq data from the cultured sample with scRNA-seq data sets from three acutely-harvested human biopsy samples confirmed that the cultures (at passage 6) express HBC markers (KRT5, TP63, EGFR) but *not* the respiratory marker SERPINB3 (**Fig. 5D**), and cluster generally with the olfactory HBCs from acute biopsies in UMAP plots (**Fig. 5E**).

**Fig. 5.**
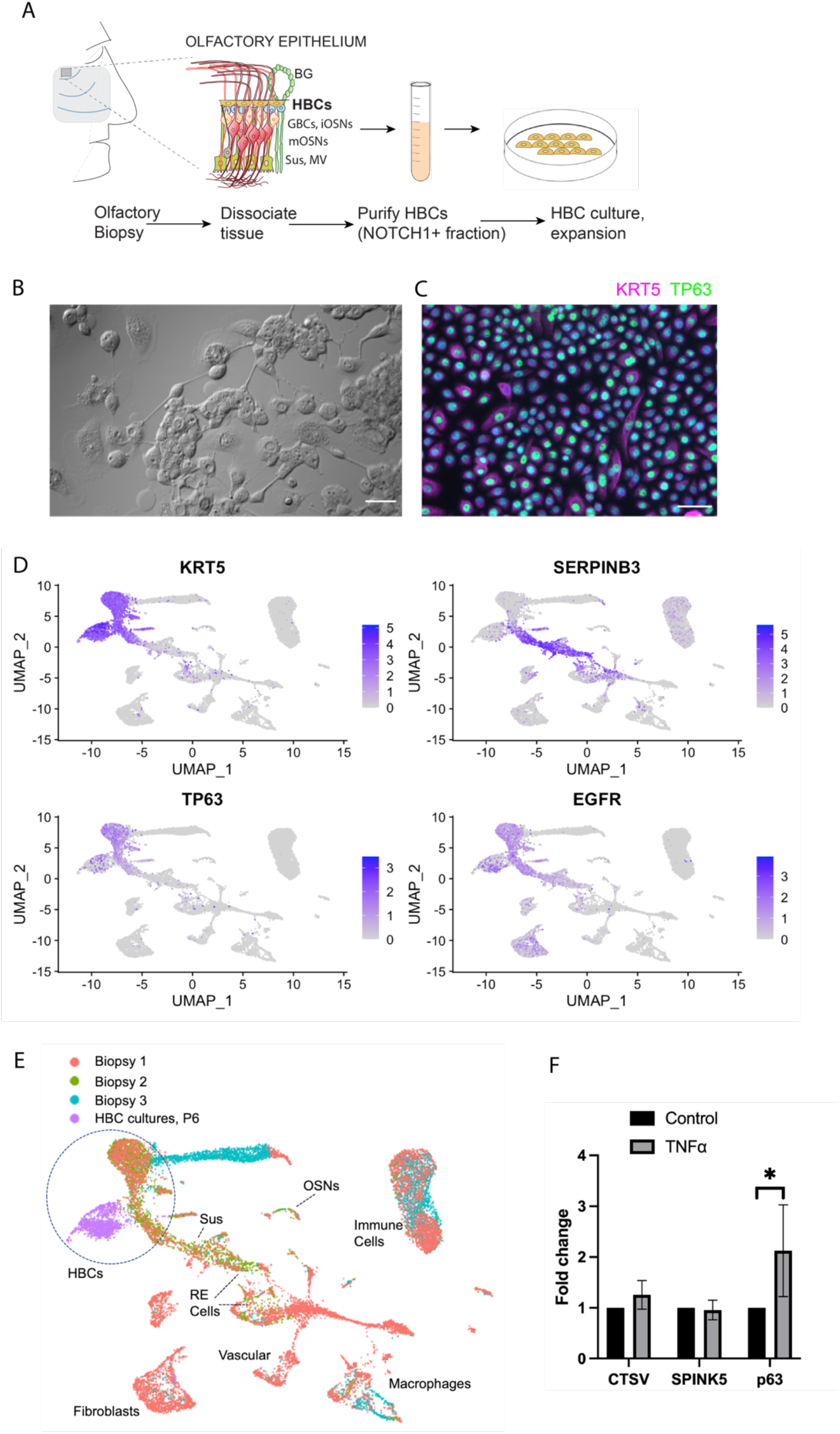
A cell culture model for human HBC analysis. **(A)** Experimental approach; human olfactory mucosa biopsy is dissociated, followed by immunoselection for NOTCH1 (+) cells, seeded onto laminin for in vitro expansion. **(B)** DIC view of typical adherent cell islands and **(C)** immunostaining of cultures showing near-uniform TP63(+)/KRT5(+) phenotype. **(D, E)** scRNA-seq analysis of Passage 6 HBC cultures; the culture single cell dataset was integrated with 3 control acutely-harvested human olfactory biopsy scRNA-seq datasets. FeaturePlots in **D** show gene expression of selected transcripts, and UMAP **(E)** showing sample origins indicates that culture-expanded cells (purple) cluster adjacent to acutely-harvested olfactory HBCs, and largely lack SERPINB3. OSNs=olfactory sensory neurons; RE=respiratory epithelium; Sus=sustentacular(**F)** RT-qPCR assay of TNFα-treated HBC cultures; note 2 fold-change increase in TP63 expression relative to controls (p=0.04, n=3). Bar=50 μM in B, C.

We next used this culture model to test the effects of TNFα on HBC gene expression. Cells were cultured on 5% laminin in either HBC growth medium with 0.5ng/mL TNFα or HBC growth medium alone for 72 hours (**Fig. 5F**). RT-qPCR assays showed no change for HBC subset markers identified in our initial scRNA-seq, however TP63 levels were increased (**Fig. 5F**). TNFα-treated HBCs exhibited more than a 2-fold increase TP63 compared to the HBCs cultured in growth medium alone (p = 0.04, t-test, n=3 replicates), suggesting that an effect of acute TNFα exposure may promote HBC quiescence, as opposed to differentiation, since TP63 acts to hold HBCs dormant. The culture model will, therefore, provide a means to further assess mechanisms impacting human HBC growth and potential pharmacologic agents.

## DISCUSSION

We provide here a single cell analysis of olfactory mucosa from presbyosmic and normosmic human subjects. Despite harboring a robust regenerative capacity, a failure of olfactory neuroepithelial maintenance or homeostasis with aging is thought to account for presbyosmia. Our results indicate that a loss of stem cells does *not* explain the histologic picture of neurogenic exhaustions that is seen in the aging human OE. Rather, we identify transcriptional alterations in HBCs, the olfactory reserve stem cells, consistent with a model in which aging-related damage and inflammation result in stem cell dysfunction (**Fig. 6**).

**Fig. 6.**
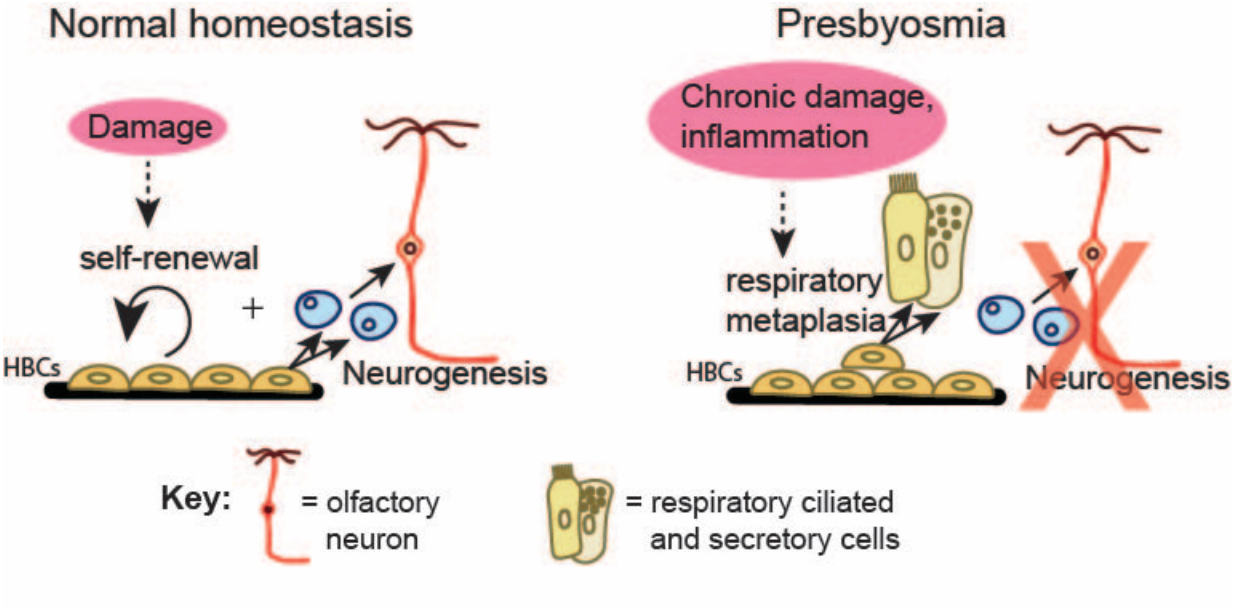
A model for presbyosmic stem cell dysfunction. Aging related damage promotes an inflammatory milieu in which olfactory HBC alterations result in decreased neurogenic activity. Histologic findings include reactive or layered HBCs along with a respiratory-like metaplasia, which might emerge from altered basal cell differentiation or from repopulation of the surface epithelium from Bowman’s gand/duct progenitors(*50*) (not depicted).

As a barrier epithelium, the immune defense functions of the nasal mucosa are well-described, especially in the setting of chronic disease states such as sinusitis or polyposis, including innate and adaptive responses (*39, 40*). How nasal immune signaling interacts specifically with the highly specialized olfactory neuroepithelial cell populations, especially the neurogenic basal cells, has only recently begun to be understood, largely in rodent models (*22*). It is evident that the OE must balance the challenges of a typical barrier epithelium with a chemosensory function requiring maintenance of intact populations of olfactory receptor neurons, and their replacement following ongoing turnover or severe damage. Unlike mouse models, the human OE must do so for many decades to avoid an age-related loss of function, i.e. presbyosmia.

Drawing on other self-renewing epithelia, such as the skin or intestinal crypts, it is clear that tissue-resident adult stem cell populations utilize similar regulatory signals, yet hold a remarkable ability to redirect their function or behavior in response to patterns of tissue injury (*41, 42*). In rodent OE, the GBCs proliferate and handle the “normal” replacement of olfactory neurons as needed, while the typically quiescent HBCs are activated by more severe damage including barrier disruption that occurs when the sustentacular population is lost (*13, 14, 16, 43*). Despite these two categories of progenitor cells, a mouse model of chronic local cytokine overexpression in the OE fails to reconstitute the neuroepithelium, resulting in HBC populations remaining abnormally dormant in the setting of neuronal and sustentacular cell loss (*22, 44*). Mechanistically, TNFα receptors on OE cells appear to mediate cytokine signaling and act via the RelA and NF-KappaB pathways in basal cells (*21*). Whether human conditions such as presbyosmia may involve inflammation-mediated stem cell dysfunction has not been determined.

Our results provide the first report combining a clinical evaluation of human subjects with presbyosmia and olfactory biopsy sampling utilizing single cell transcriptional profiling. The variability in successful capture of intact olfactory neuron populations, due to respiratory-like metaplasia, has been well-described in aging human olfactory samples, suggesting the neurogenic exhaustion hypothesis (*20*). Nonetheless, the present approach is well-suited to explore transcriptional alterations among the cell populations present in control versus presbyosmic samples. Our initial analysis of data sets suggested potential differences among HBCs in presbyosmic versus control groups, while other populations were more homogenous, motivating our further focus on the stem cell analyses. We considered various potential findings that might drive neurogenic exhaustion or respiratory-like metaplasia: (a) stem cells might be sparse or absent; (b) only respiratory-type stem or progenitors might be identifiable; or (c) olfactory stem cells might remain present but exhibit molecular alterations unique to the presbyosmic samples. Our results support the last model. Focusing attention to the HBCs, or reserve stem cells, we identified heterogeneity among this population not previously described (**Fig. 2**). Our data sets provide a basis for ongoing efforts aimed at identifying the roles of HBC-subset-specific transcripts, such as CTSV or SPINK5. The inflammatory-response phenotype identified in presbyosmic HBCs appears consistent with aging-related tissue alterations that may drive an immune-defense HBC response preventing their normal neurogenic program (**Fig. 6**). Finally, our description of a human olfactory HBC culture model will provide a platform to test in vitro modulation of HBC signaling for therapeutic potential.

Limitations of this study include the restriction of our biopsy samples to the peripheral olfactory apparatus, which is the routinely accessible portion of this sensory system. We do not exclude the possibility that presbyosmia is a heterogenous condition that may, in some subjects, be due to central sites of pathology, such as the olfactory bulbs or cortex. The analysis of olfactory mucosa biopsies is not a complete survey of the olfactory cleft, but represents a sampling of the peripheral olfactory organ. Variability in the extent and continuity of olfactory versus respiratory surface epithelium is an unavoidable reality; therefore, we avoid here direct comparisons of specific cell type yields among samples and instead focus on comparisons of the transcriptional profiles of captured cell populations.

In summary, there are currently over 54 million US residents age 65 or older (US census data) and presbyosmia is prevalent in this age group. Currently, there are no specific treatments available for sensorineural olfactory disorders, including presbyosmia. The findings and data sets provided here offer new insights into the cellular and molecular changes associated with age-related olfactory loss and should provide a resource for ongoing study. Our conclusions suggest that olfactory stem cells may be a logical therapeutic target for certain sensorineural olfactory disorders. For these findings to be applied to the clinic, pre-clinical testing of reagents targeting pathways of interest in OE stem cell cultures or animal models may facilitate pilot clinical trials.

## MATERIALS AND METHODS

### Study Design

We performed a prospective analysis of presbyosmic or control normosmic subjects, to obtain nasal mucosal biopsies for analysis. No statistical method was used to predetermine sample size; rather, we have followed current best practices for scRNA-seq studies, and, guided by goals for rare cell type identification, expected read depth, and identification of DE genes for specific clusters, utilized three subjects in each cohort (*45*). Tissue samples from a single patient were processed individually. Inclusion and exclusion criteria are discussed below, based on olfactory function scores. The research objective was to identify molecular alterations in olfactory cell populations of presbyosmics. Our initial hypothesis was that molecular alterations consistent with stem cell dysfunction would be identified. Following initiation of data analysis, our hypotheses were expanded to focus also on immune cells and immune-induced alterations in stem cells. Subjects were identified from an academic Rhinology practice, without randomization, and blinding was not feasible.

### Sample acquisition and preparation

Olfactory testing was performed using a validated assessment tool, the Smell Identification Test (Sensonics, Inc.). Biopsies of the septum from the olfactory cleft region were obtained by sinus surgeons in the Department of Head and Neck Surgery & Communication Sciences at Duke University School of Medicine via an IRB-approved protocol. Tissue was obtained from patients consented for transnasal endoscopic surgery to access the pituitary or anterior skull base. Samples for scRNA-seq were selected according to the inclusion criteria as patients presented for routine clinical care. Inclusion criteria included SIT ≥ 34 for the control group and SIT <30 and age > 65 years for the presbyosmic group. Subjects were excluded who had evidence of acute nasal or sinus inflammation or obstruction on pre-operative endoscopy and imaging. Mucosa was carefully excised from portions of the olfactory cleft along the superior nasal septum or adjacent superior medial vertical lamella of the superior turbinate, uninvolved with any pathology. Immediately after removal, samples were held on ice in Hank’s Balanced Salt Solution with 0.1% BSA and transported to the laboratory. Under a dissecting microscope, any bone or deep stroma was trimmed away from the epithelium and underlying lamina propria. A small portion of some specimens was sharply trimmed and fixed in 4% paraformaldehyde in phosphate buffered saline, cryoprotected and then snap-frozen in optimal cutting temperature medium, to be cryosectioned for histology. The remaining specimen was enzymatically dissociated using collagenase type I, dispase and DNase for approximately 30 min. Next, papain was added for 10 min, followed by 0.125% trypsin for 3 min as we have reported (*17*). Cells were filtered through a 70 μm strainer, pelleted and washed, and then treated with an erythrocyte lysis buffer and washed again. The cells were then resuspended in PBS with 0.1% bovine serum albumin and 0.2% anti-clumping agent (Invitrogen) and immediately processed for scRNA-seq using the Chromium (10X Genomics) platform. Fresh human tissue samples used to generate scRNA-seq data were exhausted in the experimental process with the exception of one sample used for cell culture (described below).

### Single cell sequencing

scRNA-seq was performed using the Chromium (10X Genomics) platform either through the Duke Molecular Genomics Core or by the Goldstein lab. Single-cell suspensions were counted using a hemocytometer, ensuring >80% viability by Trypan dye exclusion, and adjusted to 1,000 cells μl−1. Samples were run using the Chromium Single Cell 3′ Paired-End, Dual-Index Library & Gel Bead Kit v3.1 (10X Genomics). The manufacturer’s protocol was used with a target capture of 5,000-10,000 cells. Each sample was processed on an independent Chromium Single Cell G Chip; 3′ gene expression libraries were dual-index sequenced using NextSeq 150bp (Illumina) flow cells by the Duke Center for Genomic and Computational Biology Core Facility.

### Single cell RNA seq analysis

Raw base call files were analyzed using Cell Ranger v.4.0.0 using the Duke Cluster Computing platform. The ‘mkfastq’ command was used to generate FASTQ files and the ‘counts’ command was used to generate raw gene-barcode matrices aligned to the GRCh38-2020-A Ensembl 98 human genome. To combat ambient RNA contamination from cell lysis during the sample dissociation and processing steps, we utilized the SoupX R package (https://github.com/constantAmateur/SoupX) for background correction of each individual sample prior to dataset integration. The data from 6 samples was analyzed in R v.4.1.0 using the Seurat package v.4.0.3. To ensure our analysis was on high-quality cells, filtering was conducted by retaining cells that expressed 100–8,000 genes inclusive and had mitochondrial gene content <10% (*46*). Samples were normalized, scaled, and integrated as we have described (*17*). Data for all six samples, totaling 36,091 cells, were combined using the standard Seurat integration workflow for normalization and batch correction (https://satijalab.org/seurat/articles/integration_introduction.html). We chose to use 50 principal components based on results from analysis using elbow plots of standard deviation. The DimPlot() function was used to generate the uniform manifold approximation projection (UMAP) plots. Clustering was conducted using the FindNeighbors() and FindClusters() functions with a resolution parameter set to 1.0. The resulting 38 Louvain clusters were visualized in a two-dimensional UMAP representation and were annotated as described (*17*). For further analysis of cell types of interest, the subset() command was used. A new analysis on each subset was performed using the FindVariableFeatures(), ScaleData(), RunPCA() and RunUMAP() functions. New UMAP plots were generated for each subpopulation, along with FeaturePlots or ViolinPlots for specific gene expression visualization. Differential gene expression between control and experimental groups was identified using the FindMarkers() command, and we removed uninformative transcripts such as XIST or HBB from further analysis. To visualize differentially expressed genes between groups, volcano plots were generated using the ggplot2 R package to plot log fold change.

### Gene set enrichment analysis (GSEA)

The top 50 enriched genes or all genes with a p-value < 0.05 from the FindMarkers() function were input into the ToppGene suite (*47*) for gene ontology and biological pathway annotation, with Bonferroni corrected for significance testing set at p < 0.05.

### Immunohistochemistry

Cryosections were prepared from nasal epithelium biopsies as described (*17*). Briefly, after PBS rinse and any required pretreatments, tissue sections were incubated in blocking solution with 5% donkey serum, and 0.1% Triton X-100 for 30 min at room temperature. Primary antibodies (**Supplemental Table 1**) were diluted in the blocking solution and incubated overnight in a humidified chamber at 4 °C. Detection by species-specific fluorescent conjugated secondary antibodies (Jackson ImmunoResearch) was performed at room temperature for 45 min. Sections were counterstained with 4,6-diamidino-2-phenylindole (DAPI) and coverslips were mounted using Vectashield (Vector Labs) for imaging, using a Leica DMi8 microscope system (Leica Microsystems). Images were processed using ImageJ software v.2.1.0/1.53c (NIH). Scale bars were applied directly from the Leica acquisition software metadata in ImageJ Tools.

### Human olfactory HBC cultures

After loading the appropriate volume of cell suspension to the 10x Chromium controller from control sample 3, remaining cell suspension was used for cell culture. To enrich for olfactory HBCs, which have been found to utilize Notch signaling (*16*), the sample was sorted using an APC-conjugated antibody to NOTCH1 (eBiosciences #17-5765-80) followed by APC magnetic selection using the EasySep kit (Stem Cell Tech) as per instructions and plated in PneumaCult medium with a SMAD inhibitor, as we have described for mouse basal cell expansion (*48*). Cells were plated on 5% laminin at 50,000 cells per well of 6 well-plate and incubated in full growth medium: PneumaCult-Ex medium (Stemcell Tech), glutamax supplement (Invitrogen), penicillin-streptomycin (Invitrogen), 10 μM Y27632, 20 ng/ml EGF, and 10 μM A83-01 (all from Stemcell Tech). Y27 was removed from medium 24 hours after initial plating. After the initial medium change, the medium was exchanged every 48 hours. Importantly, the scRNA-seq analysis of the biopsy specimen from which this culture was established revealed almost exclusively olfactory rather than respiratory populations (see Fig. 1C, 1D), suggesting minimal potential culture contamination by respiratory basal cells. Cultures were passaged splitting 1:3 with brief trypsinization when ∼80% confluent. Cells were frozen in 90% fetal bovine serum (Invitrogen) with 10% dimethyl sulfoxide and stored in liquid nitrogen. Cell phenotype was characterized as described below.

### TNFα-treatment

Passage 4 human HBCs, cultured as described above, were plated at a concentration of 20,000 cells per well of a laminin-coated 24 well plate. TNF alpha (0.5ng/mL final concentration in PBS, human recombinant, Stemcell Tech) was added to half of the wells, while the other wells received full growth medium only, with vehicle. Medium was changed after 48 hours. After 72 hours, RNA was isolated for analysis. All assays were performed in triplicate.

### RT-qPCR

Total RNA was isolated using column purification per protocol (Zymo Research Corp., Irvine, CA, USA). DNase I on-column digestion was performed. Reverse transcription first strand cDNA synthesis was performed using Superscript IV (Invitrogen). Taqman assays (Invitrogen) were used in reactions performed on a BioRad real time thermal cycler. Reactions were performed in triplicate and at least 3 replicates were tested per condition. Fold-change calculations were performed using the 2-ΔΔCt technique (*49*), and GAPDH expression was used as a reference.

### Statistics

No statistical method was used to predetermine sample size. For each experiment, tissue samples from a single patient were processed individually. Single-cell suspensions for each sample were processed for scRNA-seq in an independent Chromium chip (10x Genomics). For differential expression analysis in Seurat, the default two-sided nonparametric Wilcoxon rank-sum test was used with Bonferroni correction using all genes in the dataset. GSEA was performed as described. qPCR comparisons were analyzed by two-tailed t test using GraphPad Prism version 9. P < 0.05 was considered significant.

## Acknowledgements

We thank the Duke Molecular Genomics Core for assistance with single cell RNA sequencing approaches. We would like to thank Benjamin M. Goldstein for expert assistance with R.

## Author contributions

Conceptualization: BJG

Methodology: AO, KI, RAH, DJ, RG, BJG

Visualization: AO, HM, EAM, BJG

Funding acquisition: AO, BJG

Supervision: HM, BJG

Writing – original draft: AO, BJG

Writing – review & editing: AO, KI, RAH, DJ, RG, EAM, HM, BJG

## Data and materials availability

Accession numbers will be made available prior to publication.

## Supplemental Materials

List of supplemental materials:

**Supplemental Figure 1. Analysis of sensory cell clusters.**

**Supplemental Figure 2. DotPlot analysis of integrated sample.**

**Supplemental Figure 3. Analysis of immune cell clusters**.

**Supplemental Table 1. Primary Antibodies**

## References

1. J. M. Pinto, K. E. Wroblewski, D. W. Kern, L. P. Schumm, M. K. McClintock, Olfactory dysfunction predicts 5-year mortality in older adults. PloS one 9, e107541 (2014).

2. T. S. Pence, E. R. Reiter, L. J. DiNardo, R. M. Costanzo, Risk factors for hazardous events in olfactory-impaired patients. JAMA Otolaryngol Head Neck Surg 140, 951–955 (2014).

3. B. Gopinath, K. J. Anstey, A. Kifley, P. Mitchell, Olfactory impairment is associated with functional disability and reduced independence among older adults. Maturitas 72, 50–55 (2012).

4. R. Choi, B. J. Goldstein, Olfactory epithelium: Cells, clinical disorders, and insights from an adult stem cell niche. Laryngoscope Investig Otolaryngol 3, 35–42 (2018).

5. K. W. Cooper, D. H. Brann, M. C. Farruggia, S. Bhutani, R. Pellegrino, T. Tsukahara, C. Weinreb, P. V. Joseph, E. D. Larson, V. Parma, M. W. Albers, L. A. Barlow, S. R. Datta Di Pizio, COVID-19 and the Chemical Senses: Supporting Players Take Center Stage. Neuron 107, 219–233 (2020).

6. V. Parma, K. Ohla, M. G. Veldhuizen, M. Y. Niv, C. E. Kelly, A. J. Bakke, K. W. Cooper, C. Bouysset, N. Pirastu, M. Dibattista, R. Kaur, M. T. Liuzza, M. Y. Pepino, V. Schopf, V. Pereda-Loth, S. B. Olsson, R. C. Gerkin, P. Rohlfs Dominguez, J. Albayay, C. Farruggia, S. Bhutani, A. W. Fjaeldstad, R. Kumar, A. Menini, M. Bensafi, M. Sandell, I. Konstantinidis, A. Di Pizio, F. Genovese, L. Ozturk, T. Thomas-Danguin, J. Frasnelli, S. Boesveldt, O. Saatci, L. R. Saraiva, C. Lin, J. Golebiowski, L. D. Hwang, M. H. Ozdener, M. D. Guardia, C. Laudamiel, M. Ritchie, J. Havlicek, D. Pierron, E. Roura, M. Navarro, A. A. Nolden, J. Lim, K. L. Whitcroft, L. R. Colquitt, C. Ferdenzi, E. V. Brindha, A. Altundag, A. Macchi, A. Nunez-Parra, Z. M. Patel, S. Fiorucci, C. M. Philpott, B. C. Smith, J. N. Lundstrom, C. Mucignat, J. K. Parker, M. van den Brink, M. Schmuker, F. P. S. Fischmeister, T. Heinbockel, V. D. C. Shields, F. Faraji, E. Santamaria, W. E. A. Fredborg, G. Morini, J. K. Olofsson, M. Jalessi, N. Karni, A. D’Errico, R. Alizadeh, R. Pellegrino, P. Meyer, C. Huart, B. Chen, G. M. Soler, M. K. Alwashahi, A. Welge-Lussen, J. Freiherr, J. H. B. de Groot, H. Klein, M. Okamoto, P. B. Singh, J. W. Hsieh, G. G. Author, D. R. Reed, T. Hummel, S. D. Munger, J. E. Hayes, More Than Smell-COVID-19 Is Associated With Severe Impairment of Smell, Taste, and Chemesthesis. Chemical senses 45, 609–622 (2020).

7. R. L. Doty, P. Shaman, S. L. Applebaum, R. Giberson, L. Siksorski, L. Rosenberg, Smell identification ability: changes with age. Science 226, 1441–1443 (1984).

8. C. Murphy, C. R. Schubert, K. J. Cruickshanks, B. E. Klein, R. Klein, D. M. Nondahl, Prevalence of olfactory impairment in older adults. JAMA : the journal of the American Medical Association 288, 2307–2312 (2002).

9. G. A. Graziadei, P. P. Graziadei, Neurogenesis and neuron regeneration in the olfactory system of mammals. II. Degeneration and reconstitution of the olfactory sensory neurons after axotomy. Journal of neurocytology 8, 197–213 (1979).

10. J. E. Schwob, W. Jang, E. H. Holbrook, B. Lin, D. B. Herrick, J. N. Peterson, J. Hewitt Coleman, Stem and progenitor cells of the mammalian olfactory epithelium: Taking poietic license. The Journal of comparative neurology 525, 1034–1054 (2017).

11. V. M. Carr, A. I. Farbman, The dynamics of cell death in the olfactory epithelium. Experimental neurology 124, 308–314 (1993).

12. M. Caggiano, J. S. Kauer, D. D. Hunter, Globose basal cells are neuronal progenitors in the olfactory epithelium: a lineage analysis using a replication-incompetent retrovirus. Neuron 13, 339–352 (1994).

13. C. T. Leung, P. A. Coulombe, R. R. Reed, Contribution of olfactory neural stem cells to tissue maintenance and regeneration. Nature neuroscience 10, 720–726 (2007).

14. R. B. Fletcher, M. S. Prasol, J. Estrada, A. Baudhuin, K. Vranizan, Y. G. Choi, J. Ngai, p63 regulates olfactory stem cell self-renewal and differentiation. Neuron 72, 748–759 (2011).

15. L. Gadye, D. Das, M. A. Sanchez, K. Street, A. Baudhuin, A. Wagner, M. B. Cole, Y. G. Choi, N. Yosef, E. Purdom, S. Dudoit, D. Risso, J. Ngai, R. B. Fletcher, Injury Activates Transient Olfactory Stem Cell States with Diverse Lineage Capacities. Cell stem cell 21, 775–790 e779 (2017).

16. D. B. Herrick, B. Lin, J. Peterson, N. Schnittke, J. E. Schwob, Notch1 maintains dormancy of olfactory horizontal basal cells, a reserve neural stem cell. Proceedings of the National Academy of Sciences of the United States of America 114, E5589–E5598 (2017).

17. M. A. Durante, S. Kurtenbach, Z. B. Sargi, J. W. Harbour, R. Choi, S. Kurtenbach, G. M. Goss, H. Matsunami, B. J. Goldstein, Single-cell analysis of olfactory neurogenesis and differentiation in adult humans. Nat Neurosci 23, 323–326 (2020).

18. E. H. Holbrook, E. Wu, W. T. Curry, D. T. Lin, J. E. Schwob, Immunohistochemical characterization of human olfactory tissue. Laryngoscope 121, 1687–1701 (2011).

19. A. S. Mobley, D. J. Rodriguez-Gil, F. Imamura, C. A. Greer, Aging in the olfactory system. Trends Neurosci 37, 77–84 (2014).

20. S. I. Paik, M. N. Lehman, A. M. Seiden, H. J. Duncan, D. V. Smith, Human olfactory biopsy. The influence of age and receptor distribution. Archives of otolaryngology--head & neck surgery 118, 731–738 (1992).

21. M. Chen, R. R. Reed, A. P. Lane, Acute inflammation regulates neuroregeneration through the NF-kappaB pathway in olfactory epithelium. Proceedings of the National Academy of Sciences of the United States of America 114, 8089–8094 (2017).

22. M. Chen, R. R. Reed, A. P. Lane, Chronic Inflammation Directs an Olfactory Stem Cell Functional Switch from Neuroregeneration to Immune Defense. Cell stem cell 25, 501–513 e505 (2019).

23. D. H. Brann, T. Tsukahara, C. Weinreb, M. Lipovsek, K. Van den Berge, B. Gong, R. Chance, I. C. Macaulay, H. J. Chou, R. B. Fletcher, D. Das, K. Street, H. R. de Bezieux, Y. G. Choi, D. Risso, S. Dudoit, E. Purdom, J. Mill, R. A. Hachem, H. Matsunami, D. W. Logan, B. J. Goldstein, M. S. Grubb, J. Ngai, S. R. Datta, Non-neuronal expression of SARS-CoV-2 entry genes in the olfactory system suggests mechanisms underlying COVID-19-associated anosmia. Sci Adv 6, (2020).

24. M. Deprez, L. E. Zaragosi, M. Truchi, C. Becavin, S. Ruiz Garcia, M. J. Arguel, M. Plaisant, V. Magnone, K. Lebrigand, S. Abelanet, F. Brau, A. Paquet, D. Pe’er, C. H. Marquette, S. Leroy, P. Barbry, A Single-Cell Atlas of the Human Healthy Airways. Am J Respir Crit Care Med 202, 1636–1645 (2020).

25. K. M. Child, D. B. Herrick, J. E. Schwob, E. H. Holbrook, W. Jang, The Neuroregenerative Capacity of Olfactory Stem Cells Is Not Limitless: Implications for Aging. The Journal of neuroscience : the official journal of the Society for Neuroscience 38, 6806–6824 (2018).

26. J. E. Schwob, S. L. Youngentob, R. C. Mezza, Reconstitution of the rat olfactory epithelium after methyl bromide-induced lesion. The Journal of comparative neurology 359, 15–37 (1995).

27. V. Turk, B. Turk, D. Turk, Lysosomal cysteine proteases: facts and opportunities. The EMBO journal 20, 4629–4633 (2001).

28. M. Siklos, M. BenAissa, G. R. Thatcher, Cysteine proteases as therapeutic targets: does selectivity matter? A systematic review of calpain and cathepsin inhibitors. Acta Pharm Sin B 5, 506–519 (2015).

29. T. Yang, D. Liang, P. J. Koch, D. Hohl, F. Kheradmand, P. A. Overbeek, Epidermal detachment, desmosomal dissociation, and destabilization of corneodesmosin in Spink5-/-mice. Genes & development 18, 2354–2358 (2004).

30. Q. Wang, Q. Lv, H. Bian, L. Yang, K. L. Guo, S. S. Ye, X. F. Dong, L. L. Tao, A novel tumor suppressor SPINK5 targets Wnt/beta-catenin signaling pathway in esophageal cancer. Cancer Med 8, 2360–2371 (2019).

31. L. M. Sevilla, V. Latorre, E. Carceller, J. Boix, D. Vodák, I. G. Mills, P. Pérez, Glucocorticoid receptor and Klf4 co-regulate anti-inflammatory genes in keratinocytes. Mol Cell Endocrinol 412, 281–289 (2015).

32. I. F. Ueki, G. Min-Oo, A. Kalinowski, E. Ballon-Landa, L. L. Lanier, J. A. Nadel, J. L. Koff, Respiratory virus-induced EGFR activation suppresses IRF1-dependent interferon lambda and antiviral defense in airway epithelium. J Exp Med 210, 1929–1936 (2013).

33. C. B. Del Debbio, M. F. Santos, C. Y. Yan, I. Ahmad, D. E. Hamassaki, Rho GTPases control ciliary epithelium cells proliferation and progenitor profile induction in vivo. Invest Ophthalmol Vis Sci 55, 2631–2641 (2014).

34. C. Berasain, M. A. Avila, Amphiregulin. Semin Cell Dev Biol 28, 31–41 (2014).

35. J. H. Ko, H. J. Kim, H. J. Jeong, H. J. Lee, J. Y. Oh, Mesenchymal Stem and Stromal Cells Harness Macrophage-Derived Amphiregulin to Maintain Tissue Homeostasis. Cell reports 30, 3806–3820 e3806 (2020).

36. L. A. Monticelli, G. F. Sonnenberg, M. C. Abt, T. Alenghat, C. G. Ziegler, T. A. Doering, J. M. Angelosanto, B. J. Laidlaw, C. Y. Yang, T. Sathaliyawala, M. Kubota, D. Turner, J. M. Diamond, A. W. Goldrath, D. L. Farber, R. G. Collman, E. J. Wherry, D. Artis, Innate lymphoid cells promote lung-tissue homeostasis after infection with influenza virus. Nat Immunol 12, 1045–1054 (2011).

37. M. Gury-BenAri, C. A. Thaiss, N. Serafini, D. R. Winter, A. Giladi, D. Lara-Astiaso, M. Levy, T. M. Salame, A. Weiner, E. David, H. Shapiro, M. Dori-Bachash, M. Pevsner-Fischer, E. Lorenzo-Vivas, H. Keren-Shaul, F. Paul, A. Harmelin, G. Eberl, S. Itzkovitz, A. Tanay, J. P. Di Santo, E. Elinav, I. Amit, The Spectrum and Regulatory Landscape of Intestinal Innate Lymphoid Cells Are Shaped by the Microbiome. Cell 166, 1231–1246 e1213 (2016).

38. N. Satoh-Takayama, T. Kato, Y. Motomura, T. Kageyama, N. Taguchi-Atarashi, R. Kinoshita-Daitoku, E. Kuroda, J. P. Di Santo, H. Mimuro, K. Moro, H. Ohno, Bacteria-Induced Group 2 Innate Lymphoid Cells in the Stomach Provide Immune Protection through Induction of IgA. Immunity 52, 635–649 e634 (2020).

39. C. Bachert, B. Marple, R. J. Schlosser, C. Hopkins, R. P. Schleimer, B. N. Lambrecht, B. M. Broker, T. Laidlaw, W. J. Song, Adult chronic rhinosinusitis. Nat Rev Dis Primers 6, 86 (2020).

40. R. C. Kern, D. B. Conley, W. Walsh, R. Chandra, A. Kato, A. Tripathi-Peters, L. C. Grammer, R. P. Schleimer, Perspectives on the etiology of chronic rhinosinusitis: an immune barrier hypothesis. Am J Rhinol 22, 549–559 (2008).

41. J. M. Wells, F. M. Watt, Diverse mechanisms for endogenous regeneration and repair in mammalian organs. Nature 557, 322–328 (2018).

42. H. Clevers, F. M. Watt, Defining Adult Stem Cells by Function, not by Phenotype. Annu Rev Biochem 87, 1015–1027 (2018).

43. R. B. Fletcher, D. Das, L. Gadye, K. N. Street, A. Baudhuin, A. Wagner, M. B. Cole, Q. Flores, Y. G. Choi, N. Yosef, E. Purdom, S. Dudoit, D. Risso, J. Ngai, Deconstructing Olfactory Stem Cell Trajectories at Single-Cell Resolution. Cell stem cell 20, 817–830 e818 (2017).

44. A. P. Lane, J. Turner, L. May, R. Reed, A genetic model of chronic rhinosinusitis-associated olfactory inflammation reveals reversible functional impairment and dramatic neuroepithelial reorganization. The Journal of neuroscience : the official journal of the Society for Neuroscience 30, 2324–2329 (2010).

45. C. C. Schmid KT, Bottcher A, Lickert H, Binder EB, Theis FJ, Heinig M, Design and power analysis for multi-sample single cell genomics experiments. bioRxiv, (2020).

46. M. D. Luecken, F. J. Theis, Current best practices in single-cell RNA-seq analysis: a tutorial. Mol Syst Biol 15, e8746 (2019).

47. J. Chen, E. E. Bardes, B. J. Aronow, A. G. Jegga, ToppGene Suite for gene list enrichment analysis and candidate gene prioritization. Nucleic Acids Res 37, W305–311 (2009).

48. B. J. Goldstein, G. M. Goss, R. Choi, D. Saur, B. Seidler, J. M. Hare, N. Chaudhari, Contribution of Polycomb group proteins to olfactory basal stem cell self-renewal in a novel c-KIT+ culture model and in vivo. Development 143, 4394–4404 (2016).

49. K. J. Livak, T. D. Schmittgen, Analysis of relative gene expression data using real-time quantitative PCR and the 2(-Delta Delta C(T)) Method. Methods 25, 402–408 (2001).

50. W. Jang, S. L. Youngentob, J. E. Schwob, Globose basal cells are required for reconstitution of olfactory epithelium after methyl bromide lesion. The Journal of comparative neurology 460, 123–140 (2003).

