## Supplemental figures for "Aging-related olfactory loss is associated with olfactory stem cell transcriptional alterations in humans"

SUPPLEMENTAL MATERIAL

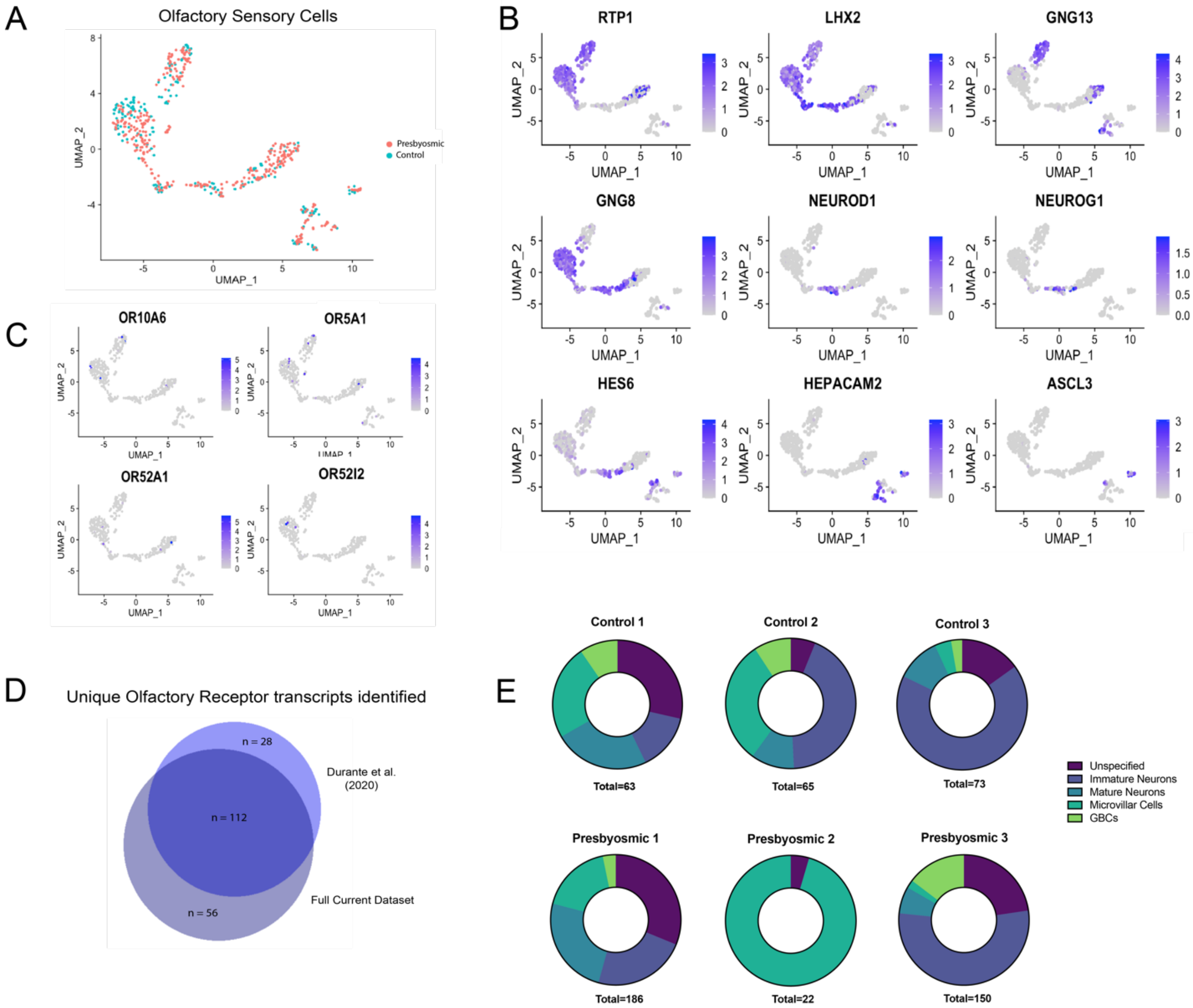


**Supplemental Figure 1. Analysis of sensory cell clusters. (A)** Sensory cell cluster comprised by mature olfactory neurons, immature olfactory neurons, GBCs and microvillar sensory cells was re-plotted as UMAP, visualized by original sample identification. **(B)** FeaturePlot visualization showing selected cell type-specific marker expression among sensory cell populations. RTP1 and LHX2 are expressed in olfactory neurons; GNG13 and GNG8 are expressed by mature or immature neurons, respectively; NEUROD1, NEUROG1 and HES6 are expressed by GBCs; HEPACAM2 and ASCL3 are expressed by microvillar sensory cells. **(C)**

FeaturePlot visualization of selected OR expression. OR10A6 and OR5A1 are Class II ORs; OR52A1 and OR 52I2 are Class I ORs; note scattered expression among neuron cell clusters, as expected. **(D)** Venn diagram depicting the unique ORs identified in our previous scRNA-seq data (*17*) and the current dataset. We identified 56 additional ORs in the present study, for a total of 196 unique OR transcripts from human biopsies of 10 subjects. See "Supplemental Table 2" spreadsheet for OR list. **(E)** Cell type composition of sensory clusters by sample. Note "Presbyosmic 2", which lacks olfactory neurons, was anosmic by SIT.

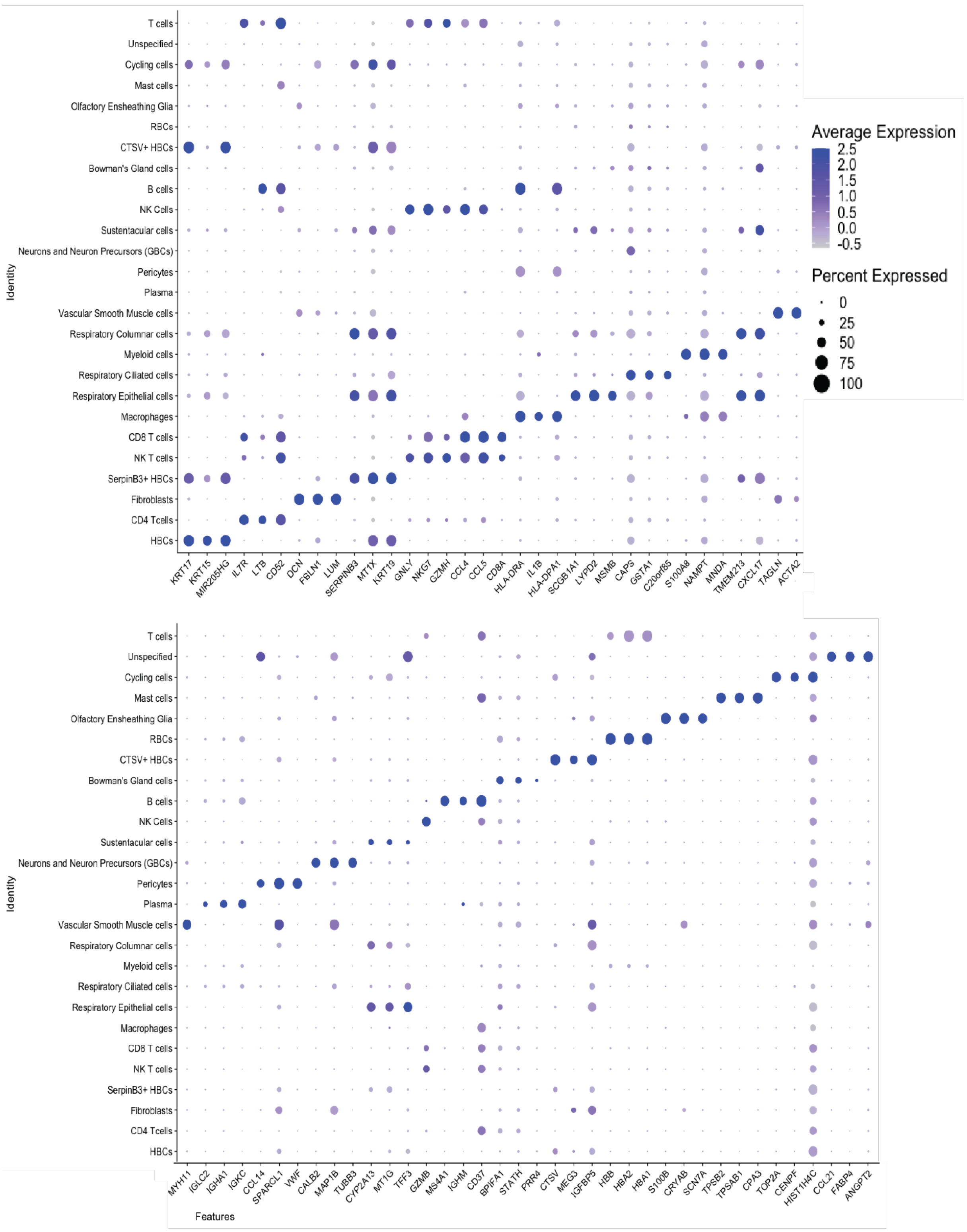


**Supplemental Figure 2. DotPlot analysis of integrated sample.** Plot depicts unbiased gene expression for top 3 enriched genes per cluster, UMAP plot is shown in Fig. 1D. Cell cluster identity is indicated on the y-axis; gene names (Features) are indicated on the x-axis. The plot depicts clusters from 36,091 cells, n=6 subjects.

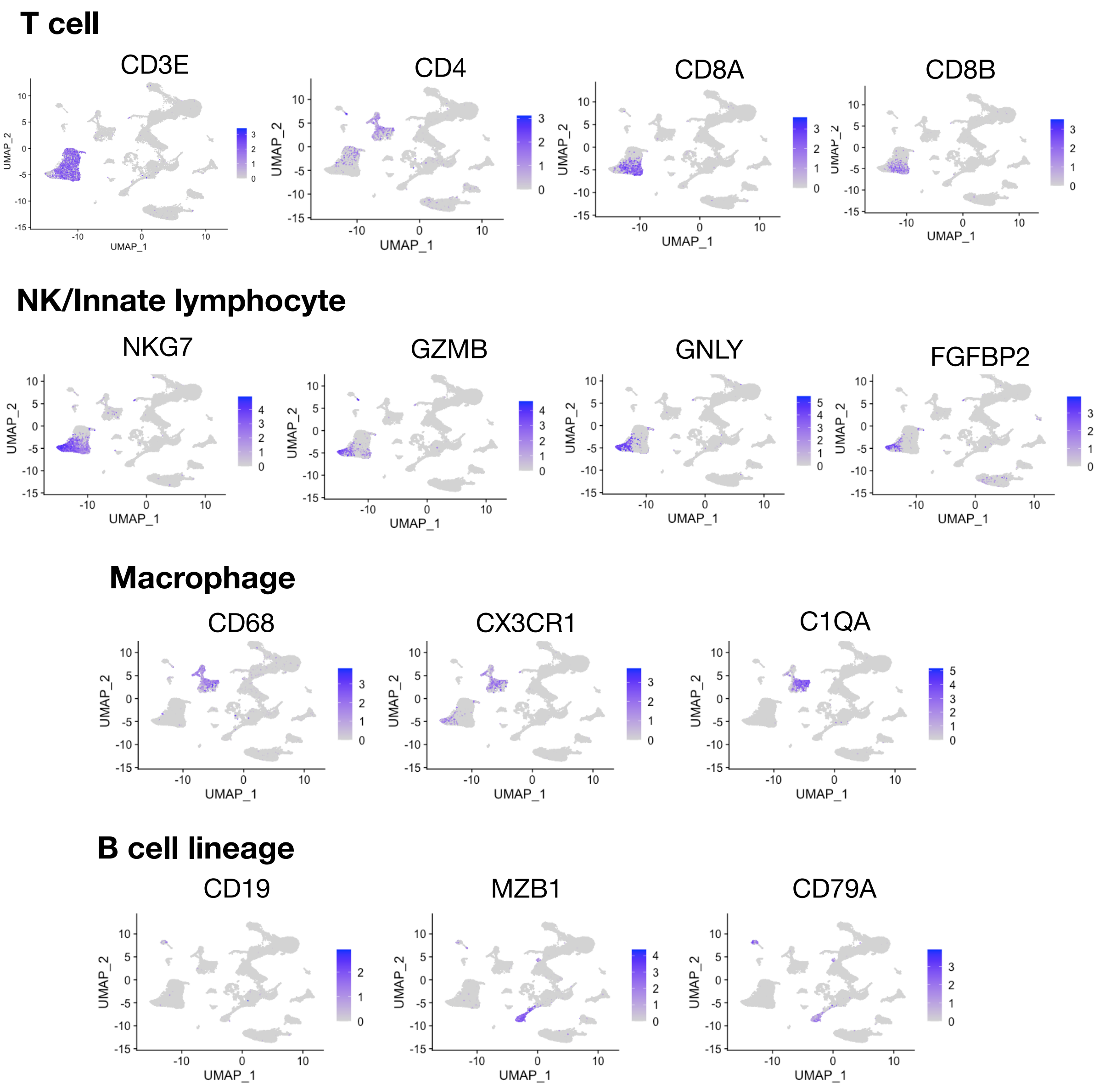


**Supplemental Figure 3. Analysis of immune cell clusters.** FeaturePlots showing indicated gene expression across integrated UMAP projections of combined data sets from normosmic and presbyosmic samples. Markers expressed by indicated immune cell phenotypes are shown, including T cells, NK/Innate lymphoid compartment (ILC) cells, Macrophages, and B cell lineages.

**Supplemental Table 1. Primary Antibodies**

| Target | Host Species | Source | Catalog Number | RRID | Dilution |
|---|---|---|---|---|---|
| CK5 | Rabbit | Abcam | ab52635 | AB_869890 | 1:1000 |
| DCX | Rabbit | Cell Signaling | 4604 | AB_561007 | 1:200 |
| SERPINB3 | Mouse | Abcam | ab180396 | AB_2892671 | 1:100 |
| SERPINB3 | Rabbit | Sigma-Aldrich | HPA055992 | AB_2682998 | 1:100 |
| TUJ1 | Mouse | Biolegend | 801201 | AB_2313773 | 1:500 |
| TUBB4 | Mouse | Sigma-Aldrich | T6793 | AB_477585 | 1:150 |
| TP63 | Mouse | Santa Cruz Biotechnology | Sc-5301 | AB_628093 | 1:500 |
